# Biomechanics and Myofibrillar Alignment Enhance Contractile Development and Reproducibility in Stem Cell Derived Cardiac Muscle

**DOI:** 10.1101/2021.05.23.445330

**Authors:** Yao-Chang Tsan, Yan-Ting Zhao, Samuel J. DePalma, Adela Capilnasiu, Yu-Wei Wu, Brynn Elder, Isabella Panse, Sabrina Friedline, Thomas S. O’Leary, Nadab Wubshet, Kenneth K. Y. Ho, Michael J. Previs, David Nordsletten, Brendon M. Baker, Lori L. Isom, Allen P. Liu, Adam S. Helms

## Abstract

Human pluripotent stem cell derived cardiomyocytes (hPSC-CMs) allow novel investigations of human cardiac disease, but disorganized mechanics and immaturity of hPSC-CMs on two-dimensional (2D) surfaces have been hurdles for efficient and reproducible study of these cells. Here, we developed a platform of micron-scale 2D cardiac tissues (M2DCTs) to precisely control biomechanics in arrays of thousands of purified, independently contracting cardiac muscle strips in 2D. By defining geometry and workload in M2DCTs in this reductionist platform that does not incorporate other cell types, we show that myofibrillar alignment and auxotonic contractions at physiologic workload critically drive maturation of cardiac contractile function, calcium handling, and electrophysiology. Additionally, the organized biomechanics in this system facilitates rapid and automated extraction of contractile kinetic parameters from brightfield microscopy images, increasing the reproducibility and throughput of pharmacologic testing. Finally, we show that M2DCTs enable precise and efficient dissection of contractile kinetics in cardiomyopathy disease models.

## Introduction

Modern techniques allow efficient generation of cardiac muscle cells from human pluripotent stem cells (hPSC-CMs), and studies of these cells have enabled new insights into human cardiac development and disease.^1–3^ However, reproducible assessment of contractile function, the most fundamental property of heart muscle, has remained a major challenge due to the immaturity of hPSC-CMs on 2D substrates, where they innately exhibit reduced membrane voltages, disorganized myofibrils, and weak contractile forces.^4,5^ For example, motion analysis and/or sarcomere shortening in routinely cultured hPSC-CMs suffers from extensive heterogeneity due to the lack of myofibrillar organization.^6–8^ Single cell micropatterning controls for cell shape in 2D and enables characterization of single cell force^9,10^, but is limited by low throughput, marked variability among replicates, and a lack of intercellular junctions that contribute to normal cardiac physiology.^11^ Three-dimensional (3D) cardiac tissues suspended between elastomer posts allow physiologic uniaxial contractions and induce the highest level of iPSC-CM maturation to date.^5^ While clearly increasing maturation, other challenges to standardized analyses with 3D tissues include 1) a requirement for stromal cell admixture^12,13^ that may increase heterogeneity and batch-to-batch variation, 2) an inability to perform concurrent imaging of myofibrils due to tissue thickness that limits comparisons across groups, 3) limited throughput, and 4) requirement for expertise in 3D tissue techniques.^5^ Thus, a 2D method that could capture the improved biomechanical aspects of 3D tissues, even if not benefitting from the complex mixed-cell type environment, would enable a simplified platform for investigating contractile function in iPSC-CMs.

The primary biomechanical drivers of improved development in 3D cardiac tissues are likely the alignment and fractional shortening that occur during coordinated contractions against elastomer substrates, but dissection of individual contributions in this system is challenging due to stromal cell admixture.^5^ Here, we developed a reductionist approach to simultaneously test the importance of tissue mechanics on cardiac maturation while also improving reproducibility of contractile analyses in 2D. In our “micron-scale 2D cardiac tissue” (M2DCT) approach, we use micropatterning of elastic polydimethylsiloxane (PDMS) to anchor thin, purified cardiac tissue strips in 2D arrays with far superior attachment rate and yield than has been possible with single cells. In these arrays, individual cardiac muscle tissues have a controlled geometry and contract independently against a defined workload. Using this highly organized biomechanical environment, we show that myofibrillar alignment and physiologic (auxotonic) contractions in M2DCTs drive marked developmental improvements in contractile function, calcium handling, and electrophysiology. Moreover, using an automated contractile analysis method (“ContractQuant”), we show that the M2DCT method reduces variability across replicates, diminishes the confounding effect of myofibrillar disorganization, and increases statistical power for pharmacologic testing. Finally, we demonstrate that the reproducible contractile microenvironment of M2DCTs allows them to be used as a test bed for investigation of developmental cues that intersect with contractile function, and as a platform for precise dissection of contractile kinetics in cardiomyopathy disease models in a simplified 2D system.

## Results

### Micron-Scale, Two-Dimensional Micropatterning Enables Robust Cardiac Tissue Formation and Myofibrillar Alignment without Stromal Cells

Based on prior observations of suboptimal myofibrillar development in single micropatterned hPSC-CMs^14^, we hypothesized that a lack of connectivity with adjacent hPSC-CMs stalls developmental signals. To test this hypothesis, we compared cell size of isolated hPSC-CMs to hPSC-CMs in contact with other hPSC-CMs. Cardiomyocytes in contact with at least one other cardiomyocyte grew to significantly larger sizes than isolated cells (**Supplemental Figure 1**). This finding motivated the development of a platform to generate geometrically defined multicellular cardiac tissue constructs that harness the benefits of reproducible myofibrillar alignment and uniaxial contractility via micropatterning^9,10,14^, while also enabling hPSC-CM connectivity.

**Figure 1.**
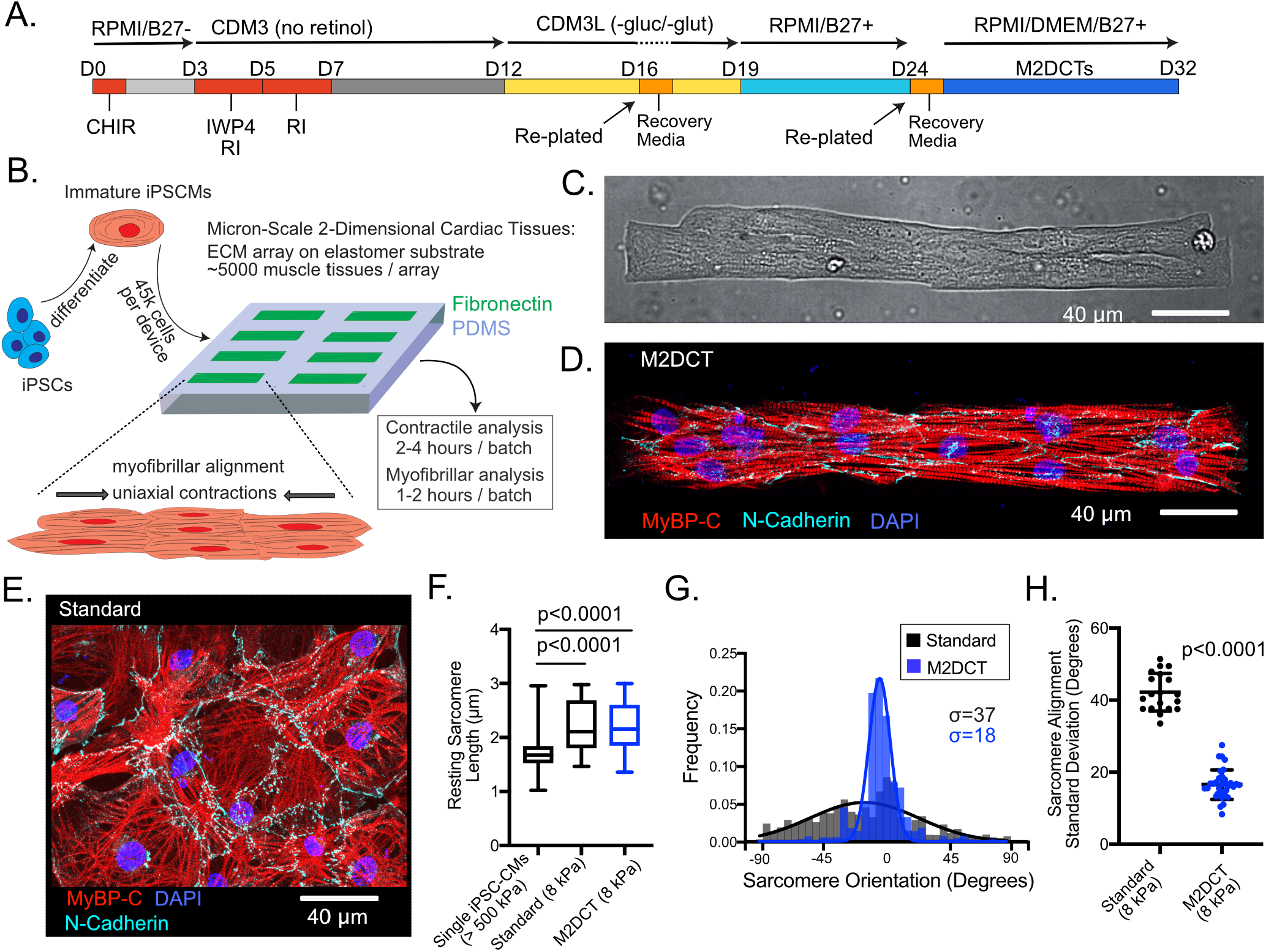
Micron-scale two-dimensional tissues (M2DCTs) exhibit enhanced myofibrillar organization. **A**. iPSCs were differentiated through wnt modulation with retinoic signaling inhibition on days 3-7. During lactate purification, CDM3 was formulated with RPMI without glucose and without glutamine to improve purification. **B**. Day 25 hPSC-CMs were replated to 8 kPa PDMS micropatterned with fibronectin as the extracellular matrix (ECM) ligand in 7:1 aspect ratio, 308 μm × 45 μm rectangles, each separated by a buffering space to minimize mechanical interactions among adjacent tissues on deformable PDMS. The overall pipeline improves efficiency by requiring only 45k hPSC-CMs per substrate, each yielding a large number of high-quality, matured M2DCTs per device for structure and function studies. The approach reduces overall time required for contractile and myofibrillar structural analyses, including both imaging and running automated image processing algorithms. **C**. hPSC-CMs avidly adhere and conform to PDMS micropatterns but not to surrounding PDMS to generate M2DCTs (image from brightfield microscopy with 40X objective). **D**. Myofibrils (marked by MyBP-C) develop along the M2DCT long-axis traversing continuously across cell junctions (marked by N-Cadherin). hPSC-CM plating density was controlled to attain an average of 6-12 hPSC-CMs per M2DCT. **E**. Representative standard (nonpatterned) hPSC-CM multicellular tissue on 8 kPa PDMS exhibits disorganized myofibrils and cell junctions in the lack of directional ECM cues from the substrate. **F**. Resting sarcomere length was measured in single micropatterned hPSC-CMs compared to non-patterned (standard) hPSC-CMs on 8 kPa PDMS and M2DCTs on 8 kPa PDMS (N=448, 229, 373 sarcomere length measurements per condition). **G**. Individual sarcomere unit orientations were measured across images using an automated imaging method in MATLAB and were fit to a Gaussian distribution for each image. A representative distribution of sarcomere orientations is shown for an M2DCT (blue) overlayed on a representative distribution for standard hPSC-CMs (black) and the standard deviation (σ) for each is shown. **H**. The standard deviation of sarcomere angles is significantly lower for M2DCTs compared to standard hPSC-CMs (N=34 and 19 tissues, respectively, with each distribution compiled from hundreds of individual sarcomere angle measurements).

Optimal induction of myofibrillar alignment and contractility in single cardiomyocytes is observed with a 7:1 aspect ratio (simulating adult myofiber eccentricity).^9,15^ Therefore, we scaled our prior single cell micropatterning approach^11,14^ by 8-fold (308 μm × 45 μm) for the M2DCT platform (**Figure 1**). The design separates individual micropatterns by a buffering distance to mechanically decouple adjacent microtissues (4,772 patterns covering 26% of the surface area of a 1.6 by 1.6 cm stamp area). We tested these micropatterning designs initially on both elastic polyacrylamide gels^14,16^ and PDMS substrates^17–19^ by microprinting fibronectin as the extracellular matrix protein. Only M2DCTs on micropatterned PDMS maintained highly consistent adherence past 3 days, and therefore elastic (~8 kPa stiffness) PDMS was subsequently utilized for all experiments. When seeded at a density to achieve 6-12 cells per micropattern at post-differentiation day 24, M2DCTs rapidly developed myofibrillar alignment and uniaxial contractions by 3 days, with consistent long-axis orientation of myofibrils across cell junctions by day 8 (**Figure 1A-D, Supplemental Figure 2, Supplemental Video 1**).

To compare myofibrillar alignment in M2DCTs and standard hPSC-CMs, we imaged myofibrils, marked by the sarcomeric protein myosin binding protein C (MyBP-C), at day 32 and quantified individual sarcomere lengths and orientations. Resting sarcomere length in both M2DCTs and non-patterned hPSC-CMs on 8 kPa PDMS substrates was ~2.2 μm (2.2 ± 0.4 μm and 2.2 ± 0.5 μm, respectively), both longer than in single hPSC-CMs on stiff PDMS (1.8 ± 0.4 μm, p<0.0001; **Figure 1E**). However, non-patterned hPSC-CMs on 8 kPa PDMS exhibited random orientations of myofibrils, in contrast to the highly reproducible alignment of myofibrils along the long axis of M2DCTs, irrespective of individual cell boundaries (**Figure 1G-J, Supplemental Figure 2**). Taken together, these results show that the M2DCT approach yields highly organized 2D purified cardiac muscle tissues with long-axis myofibrillar alignment.

### Mechanical Uncoupling of M2DCTs Results in High Fidelity Contractile Quantification with Brightfield Imaging

A major barrier to accurate quantification of hPSC-CM contractility on 2D substrates has been the heterogeneous mechanical environment caused by both myofibrillar disorganization within cells and intercellular coupling disorganization between cells. Since our goal was to mechanically decouple adjacent M2DCTs, we performed a modeling analysis to examine the influence of varying spatial relationships among M2DCTs and the PDMS substrate (**Figure 2A, Supplemental Notes**). We first analyzed effects of the PDMS buffering space surrounding each M2DCT. In an M2DCT with up to an 11% fractional shortening, we found that increasing the size of the buffer space further would result in similar fractional shortening measurements, indicating a minimal boundary effect at edges of the specified buffering spaces (**Supplemental Figure 3**). Additionally, we tested the impact of the PDMS thickness, since elastomer tethering to an underlying rigid glass surface could cause variations due to fabrication inconsistencies. We showed that 8 kPa PDMS completely uncouples contracting M2DCTs from the underlying glass at any thickness greater than 40 μm (**Supplemental Figure 3**). Since our method utilizes measurements of fractional shortening, we also assessed how the tissue elastic modulus may influence relationships between fractional shortening and force generation. In the range of cardiac tissue elastic moduli (8-12 kPa)^20^, the influence of variability in M2DCTs’ elastic moduli was minor on 8 kPa PDMS (**Supplemental Figure 3**). Finally, we modeled the relationship between fractional shortening measured at different locations within M2DCTs and whole tissue force to determine optimal locations for region of interest (ROI) placement when measuring fractional shortening. Based on these results, we established that ROI locations measuring displacements of the inner 50% of the tissue length are most robust to minor variations in ROI placement (**Supplemental Figure 3**).

**Figure 2.**
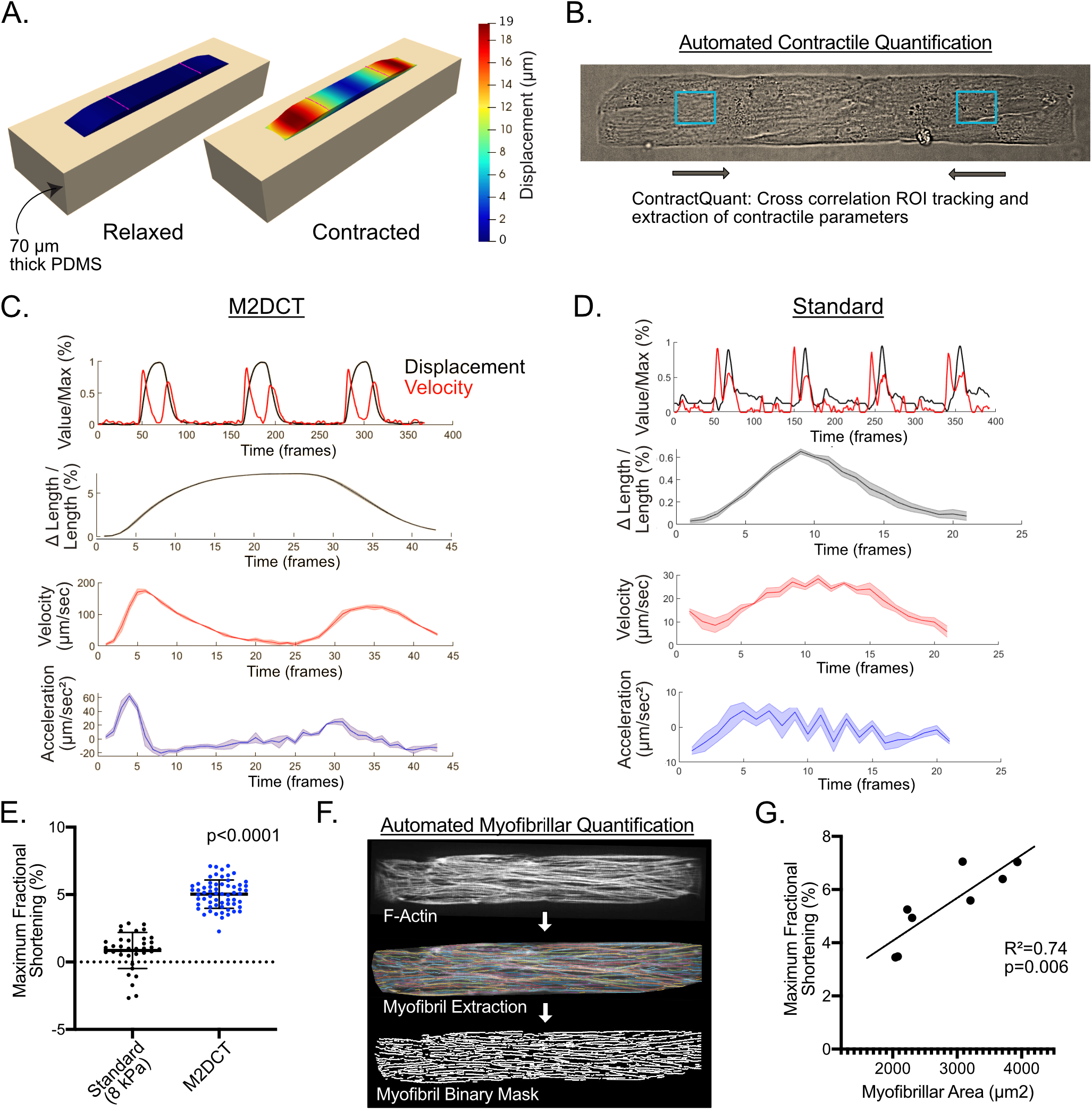
Mechanical uncoupling of adjacent M2DCTs improves contractile function and quantification reproducibility. **A**. Modeling analysis demonstrated that M2DCTs are mechanically uncoupled from adjacent M2DCTs and the glass underlying 8 kPa PDMS at the thickness and mechanical buffering spaces used. 3D representation of an M2DCT on 70 μm thick PDMS at 11% fractional shortening shown with heat map depicting local displacements. Pink lines demarcate the inner 50% length of the M2DCTs where regional tissue deformations are most stable for accurate measurements of fractional shortening. **B**. Brightfield image of M2DCT with regions of interest (ROIs) annotated for displacement tracking using cross correlation in MATLAB with the ContractQuant algorithm. **C**. Representative output from ContractQuant for an M2DCT showing (top) concurrent normalized displacements and velocities for the full time series capture, (second row) fractional shortening of merged contractions, (third row) merged absolute values of contractile velocity, and (fourth row) absolute values for acceleration. In the merged contraction data (rows 2-4), 95% confidence intervals are shown (shaded regions), indicating minimal beat-to-beat variability. **D**. Representative output from ContractQuant for standard, non-patterned hPSC-CMs on 8 kPa PDMS shows significantly greater heterogeneity within and across individual contractions that prevents extraction of parameters other than maximal fractional shortening. **E**. Maximal fractional shortening is greater in M2DCTs than standard hPSC-CMs from the same batches on 8 kPa PDMS (N=62 and 36 respectively). **F**. Myofibrils in M2DCTs were routinely imaged (SiR-actin, to label F-actin) following contractile analysis to verify cardiomyocyte purity and homogeneous distribution of myofibrils in analyzed tissues. An automated image analysis pipeline was used to extract signal peaks corresponding to individual myofibrillar bundles and then estimate total myofibrillar bundle area for each M2DCT. **G**. Batch variation was assessed by comparing mean myofibrillar area and mean maximal fractional shortening (N=150 M2DCTs across 7 batches; linear fit assessed by R^2^; comparison to slope of 0 by extra sum of square F test).

Next, we developed a method to quantify contractile function from M2DCTs. To avoid phototoxicity-induced contractile artifacts, we developed the ContractQuant algorithm, which tracks pixel features of ROIs from brightfield time series images (50 frames-per-second) using a cross-correlation method implemented in MATLAB (**Figure 2B-D**). The reproducible contractile behavior of M2DCTs enable calculation of contraction and relaxation velocities at multiple phases of each contractile cycle by displacement tracking within ROIs positioned within the inner 50% of M2DCTs (**Figure 2C**). MATLAB scripts for ContractQuant are available on Github. Taken together, we show that contractile quantification is highly tractable in M2DCTs due to the consistent long-axis uniaxial contractile behavior of individual tissues that are mechanically decoupled from each other.

### Defined Biomechanics in M2DCTs Markedly Improves Myofilament Function and Reproducibility Across Replicates

Next, we compared maximal fractional shortening, as a measure of myofilament function, in M2DCTs versus standard hPSC-CMs. Reproducible measurements of parameters other than fractional shortening in standard hPSC-CMs were not possible due to excessive irregularity in contractile behavior (**Figure 2D**). Average maximal fractional shortening was significantly greater in M2DCTs compared to standard hPSC-CMs from the same batches (5.0±1.1 vs. 0.9±1.4, p<0.0001, **Figure 2E**). Moreover, variability across M2DCTs was reduced by 87% compared to standard hPSC-CMs (coefficient of variation 0.21 vs. 1.58, **Figure 2E**). In addition, standard hPSC-CMs exhibited regions of apparent stretch due to coupling with adjacent, more strongly contracting cells (**Supplemental Video 2**). Using the modeling approach described above, we calculated the traction force generated by M2DCTs on 8 kPa PDMS; M2DCTs generated a mean traction stress of 404 μN/mm^2^.

We assessed the influence of batch to batch variability by measuring contractile function concurrently with myofibrillar structure for 150 M2DCTs across 7 differentiation batches. Taking advantage of the ease of myofibrillar imaging in thin M2DCTs, we used an automated myofibrillar detection method (“MyoQuant”) from fluorescence images of F-actin that followed contractile imaging (**Figure 2F**).^11^ We found that variability in contractile function in M2DCTs was largely due to differences in myofibrillar abundance, with a significant correlation between mean myofibrillar area per batch and mean maximal fractional shortening (R^2^=0.74, p=0.006 vs. slope of 0; **Figure 2G**). Combining this approach with the M2DCT platform also enables rapid quality control assessment, including verification of hPSC-CM purity and filtering of tissues without homogeneous distributions of myofibrils from contractile analyses (**Supplemental Figure 4**).

### Maturation of Calcium Handling, Electrophysiology, and Gap Junction Development is Dependent on the Biomechanical and Contractile Environment

Next, we examined whether the improved myofibrillar structure and function associated with the organized biomechanical environment of M2DCTs results in maturation of calcium handling, ion channel and gap junction expression (**Figure 3A**). We found that *SCN5A* and *KCNJ2* were both markedly upregulated in M2DCTs relative to standard hPSC-CMs, indicating greater expression of encoded Nav1.5 (responsible for phase 0 depolarization) and Kir2.1 (responsible for maintaining negative membrane potential), respectively. *HCN4*, responsible for sodium influx in pacemaking cells, was down-regulated in M2DCTs. *ATP2A2, RYR2*, and *CASQ2* genes were significantly up-regulated, consistent with increased expression of the sarcoplasmic reticulum proteins, SERCA2, ryanodine-2, and calsequestrin-2, respectively. Finally, the predominant cardiac gap junction gene, *GJA1* (encoding connexin-43), was markedly up-regulated in M2DCTs.

**Figure 3.**
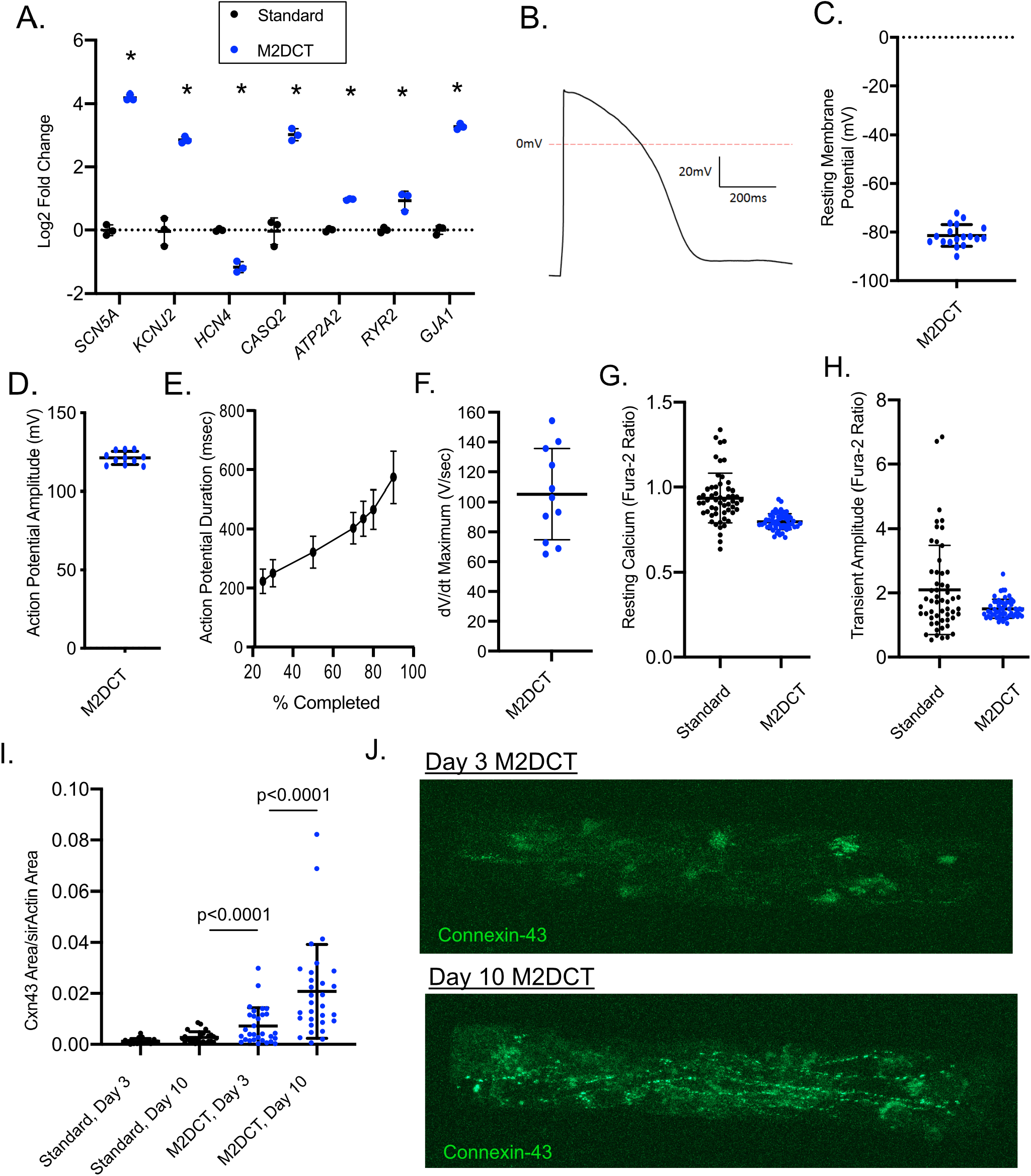
Maturation of Calcium Handling, Electrophysiology, and Gap Junction Development is Dependent on the Biomechanical and Contractile Environment. **A**. RNA expression of ion channel and calcium handling genes by RNA-seq in M2DCTs was compared to standard hPSC-CMs using RNA-seq reads normalized by DE-Seq2. **B**. Representative action potential from an M2DCT by patch clamping. **C-F**. Patch clamping of M2DCTs showed reproducible resting membrane potentials(C), action potential amplitude (D), action potential duration (E), and maximum dV/dt during phase 0 depolarization (F). Resting membrane potentials were measured from 18 M2DCTs from 5 batches and triggered action potentials were recorded for 11 M2DCTs from 4 batches. **G**. Resting fura-2 ratios were lower in M2DCTs with reduced variability across samples. **H**. Calcium transient amplitudes exhibited less variability in M2DCTs. **I**. Connexin-43 quantification by time lapse immunofluorescence (day 3 and 10) using a live cell GFP reporter line (Allen Institute AICS-0053 cl.16) demonstrated greater abundance in M2DCTs with a marked increase between days 3 and 10. **J**. Paired images of a representative M2DCT at day 3 and 10 from the connexin-43-GFP reporter line.

To determine whether these gene expression differences correlated with functional improvements, we performed patch clamping in M2DCTs. We reproducibly observed mature resting membrane potentials in M2DCTs (−81 ± 4 mV, **Figure 3B-C**), as compared to prior characterizations of immature hPSC-CMs.^5^ Analysis of triggered action potentials also revealed a more mature profile – action potential amplitude was 121 ± 4 mV; action potential duration at 90% was 578 ± 88 msec; and maximum dV/dt (phase 0 depolarization) was 105 ± 31 V/sec (**Figure 3D-F**). We assessed calcium handling with the fura-2 indicator dye. M2DCTs exhibited lower resting calcium levels than standard hPSC-CMs, indicating greater sarcoplasmic reticulum retention consistent with higher *ATP2A2* and *CASQ2* expression (**Figure 3E**). Average calcium transient amplitudes were similar in M2DCTs, but variability across tissues was 73% less in M2DCTs (standard deviation of calcium transient amplitude 0.29 vs. 1.06 fura-2 ratio, p<0.0001, **Figure 3F**). Finally, we quantified gap junction formation, which is essential for mature electromechanical coupling in heart tissue. We imaged connexin-43 using a GFP reporter line at day 3 and day 10 following M2DCT formation. M2DCTs exhibited greater connexin 43 expression at day 3 compared to standard hPSC-CMs, but with predominant perinuclear localization (**Figure 3G-H**). Paired analysis of the same M2DCTs imaged at day 10 exhibited a marked increase in connexin-43 abundance with localization at cell junctions.

### M2DCTs Increase the Precision of Pharmacologic Testing in hPSC-CMs

Pharmacologic testing is a major application of hPSC-CMs but immaturity and variability of these cells may limit its reliability. To test whether the controlled biomechanical environment of M2DCTs improves reproducibility, we quantified fractional shortening of M2DCTs in the presence of contractile antagonists and agonists. We demonstrated that the contractile inhibitors mavacamten (myosin ATPase inhibitor) and verapamil (L-type calcium channel inhibitor) both exhibited a highly reproducible dose-dependent decrement in fractional shortening relative to standard hPSC-CMs (**Figure 4A-D**). Increasing media calcium concentration elicited a reproducible dose-related increment in fractional shortening (**Figure 4E**). Quantification of contractility in response to mavacamten in M2DCTs from a separate iPSC line also showed a reproducible dose-response (**Figure 4F**). Based on the observed standard deviations, a power analysis revealed that the number of samples needed to quantify a difference of 20% in independent samples with alpha=0.05 and beta=0.1 is reduced from 205 for non-patterned hPSC-CMs to 23 using M2DCTs.

**Figure 4.**
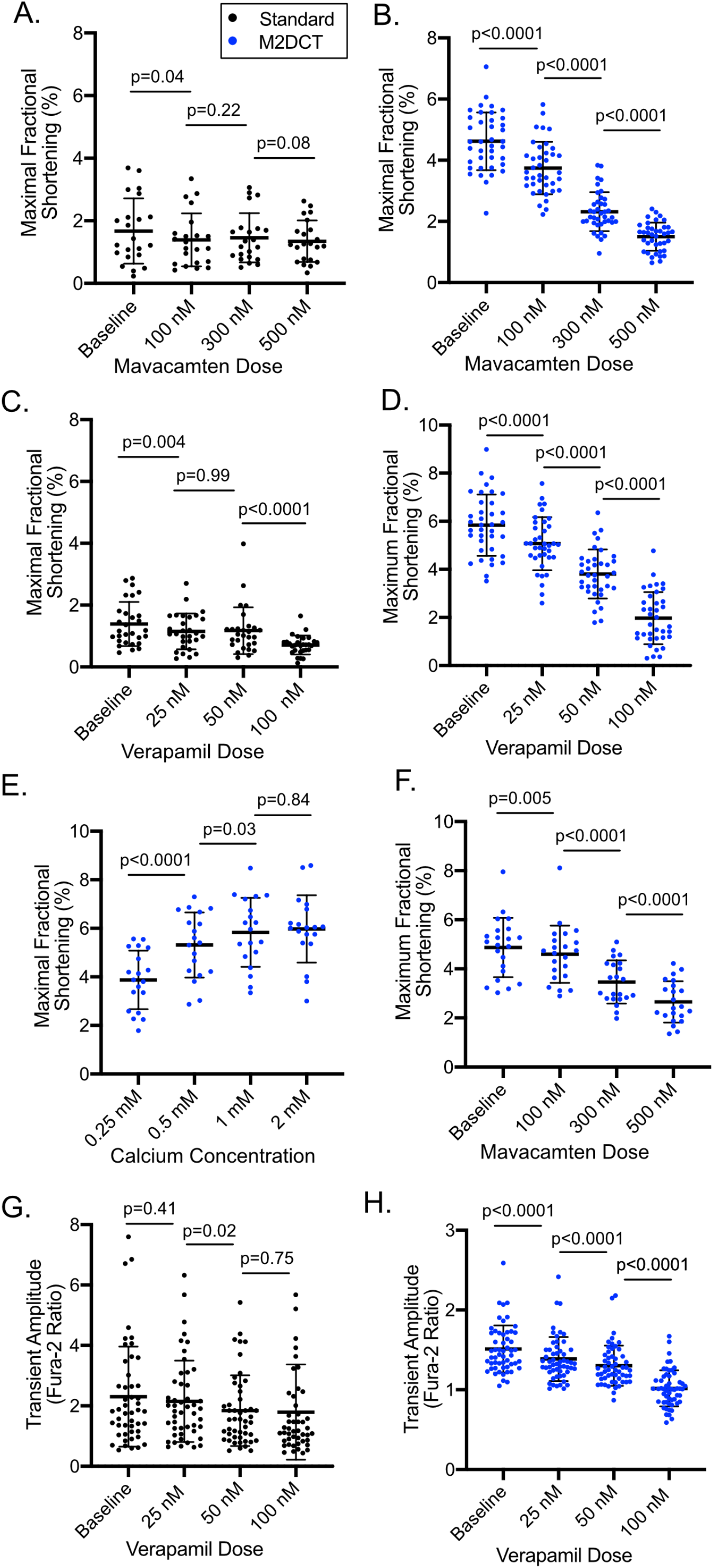
M2DCTs Increase the Precision of Pharmacologic Testing in hPSC-CMs. **A-B**. Maximal fractional shortening with verapamil in standard hPSC-CMs (A) and M2DCTs (B) on 8 kPa PDMS. **C-D**. Maximal fractional shortening with myk-461 in standard hPSC-CMs (C) and M2DCTs (D) on 8 kPa PDMS. **E**. Dose-response of increasing calcium concentration on maximal fractional shortening in M2DCTs. **F**. Maximal fractional shortening with myk-461 in M2DCTs on 8 kPa PDMS from a different iPSC line (Allen Institute AICS-0053 cl.16). **G-H**. Calcium transient amplitudes with verapamil in standard hPSC-CMs (G) and M2DCTs (H).

Additionally, we found that M2DCTs demonstrated less variability than standard hPSC-CMs in calcium transients in response to successive doses of verapamil (**Figure 4G-H**). Based on the observed standard deviations, a power analysis revealed that the number of samples needed to quantify a difference of 20% in independent samples with alpha=0.05 and beta=0.1 is reduced from 148 to 12 using M2DCTs.

### Maturation of hPSC-CMs is Induced by Signals Integrated from Biomechanical and Metabolic Inputs

Next, we compared the maturation effect of the biomechanical environment of M2DCTs to maturation achieved through the following alternate biochemical techniques: 1) oxidative phosphorylation promoting media (“OxPhos”, modified from Correia et al.^21^), 2) tri-iodothyronine thyroid hormone (T3),^22–24^ 3) T3 plus dexamethasone,^23,24^ and 4) cell replication inhibition by torin1.^25^ We first validated a maturation effect of OxPhos media in standard hPSC-CMs under our culture conditions, using the proportions of cardiac troponin I (cTnI) and slow skeletal troponin I (ssTnI) as a maturation read-out.^5,26^ OxPhos media induced an immediate increase in cTnI in monolayer hPSC-CMs, attaining 56 ± 4% of total TnI after only 5 days (**Figure 5A**). We showed that this switch occurs through differential expression of *TNNI3* mRNA (encoding cTnI) relative to *TNNI1* mRNA (encoding ssTnI), with further augmentation of *TNNI3* by addition of T3 (**Figure 5B**). Based on these results, we also included M2DCTs maintained in OxPhos-T3 media to test whether maturation in M2DCTs can be further enhanced.

**Figure 5.**
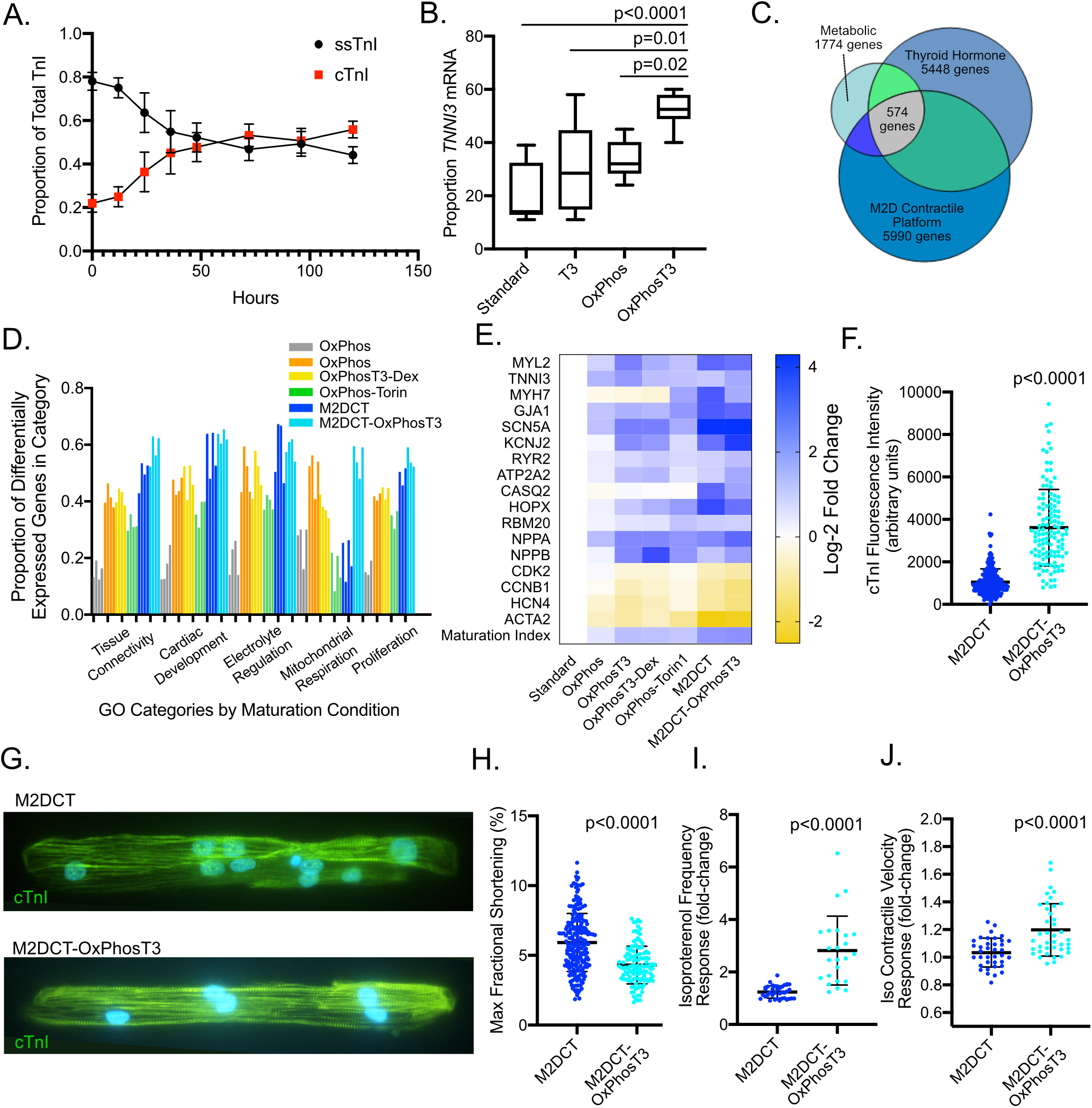
Maturation of hPSC-CMs is Induced by Interacting Signals from the Biomechanical and Metabolic Environments. **A**. In standard hPSC-CM monolayers, switching to a media that requires oxidative phosphorylation (OxPhos) induces expression of cTnI compared to ssTnI (measured by mass spectroscopy, N=6 at each time point, duplicates for 3 different hPSC-CM lines) beginning in the first 24 hours after media change. **B**. Relative *TNNI3* mRNA expression (as a proportion to total of *TNNI3* + *TNNI1* mRNA) is increased in standard hPSC-CMs in OxPhos media synergistically with T3 after 5 days (N=6, triplicates for 2 different hPSC-CM lines). **C**. Venn diagram showing overlap among differentially expressed genes (DE-seq2 adjusted p <0.001) in different maturation conditions. **D**. Gene ontology analysis of maturation conditions versus standard monolayer hPSC-CMs demonstrated over-representation of differentially expressed genes among categories associated with tissue connectivity (e.g. multicellular organism development, extracellular matrix organization, response to mechanical stimulus, cell-cell junction organization shown left-right), cardiac development (e.g. cardiac muscle cell differentiation, actin cytoskeleton organization, cardiac muscle hypertrophy, cardiac ventricle development shown left-right), electrophysiology (e.g. regulation of membrane potential, cardiac muscle cell action potential, membrane repolarization, cellular calcium ion homeostasis shown left-right), mitochondrial respiration (e.g. cellular respiration, oxidative phosphorylation, mitochondrion organization, respiratory electron transport chain shown left-right), and cell proliferation (e.g. regulation of cell migration, regulation of cell proliferation, muscle cell proliferation shown left-right). Data is shown as proportion of differentially expressed genes (DE-seq2 p<0.001) out of all genes in each category (numbers of genes and adjusted p-values for each gene sets shown are in Supplementary Table 2). **E**. Relative expression (log-2 fold change) of genes associated with maturation is shown by heat map across maturation conditions. In the bottom row, a maturation index was calculated as the average of the fold-changes across all of these genes (taking the negative value for genes inversely associated with maturation – i.e. cell cycle genes *CDK2* and *CCNB1*, sodium influx gene *HCN4*, and smooth muscle α-actin gene *ACTA2*). **F**. Quantitative immunofluorescence of cTnI in hPSC-CMs in M2DCTs in regular glucose-containing media or OxPhosT3 media. **G**. Representative images of cTnI expression in M2DCTs in regular (top) and OxPhosT3 media (bottom). More hPSC-CMs in regular media exhibited minimal myofibrillar expression of cTnI (lower cells in top image). **H**. Contractile quantification in M2DCTs in standard and OxPhosT3 media. **I**. Isoproterenol frequency response (10 nM) was quantified as fold change from baseline for each tissue. **J**. Response of M2DCT contraction velocity to isoproterenol 10 nM was quantified as fold change from baseline for each tissue.

Comparison of differentially regulated genes (adjusted p<0.001) revealed both shared and distinct differentially regulated genes among different maturation strategies (**Figure 5C**). Sample groups clustered separately in a principal component analysis, while housekeeping genes^27^ showed similar expression across conditions, verifying robust normalization (**Supplemental Figures 5A-B**). In M2DCTs, gene ontology (GO) showed over-enrichment of 561 biologic process categories (**Supplemental Table 1**). Many GO categories across maturation conditions contained overlapping genes and could be summarized in 5 general types – tissue connectivity, cardiac development, electrophysiology, mitochondrial respiration, and cell proliferation (**Figure 5D, Supplemental Table 2)**. While most categories exhibited a larger extent of differential expression in M2DCTs, an exception was mitochondrial respiration. Genes in these categories were not significantly differentially expressed in M2DCTs but exhibited a markedly different profile in M2DCTs in OxPhos-T3 media.

A panel of genes strongly associated with maturation in hPSC-CMs^5^ revealed improved maturation with all of the tested strategies. However, using a transcriptomic maturation index, M2DCT conditions exhibited the largest overall gain (**Figure 5E**). Notably, *HOPX*, which was recently shown to be an important regulator of growth and maturation in hPSC-CMs^28^, was increased by 12-fold in M2DCTs.

Although overall differential expression profiles were generally similar between M2DCTs in regular media versus OxPhos-T3, a larger *TNNI3* increase was present in the OxPhosT3 condition. We assayed differential cTnI expression in M2DCTs in OxPhos-T3 media using quantitative immunofluorescence. The mean abundance of cTnI was greater in OxPhos-T3 M2DCTs (3608 ± 1801 vs. 1044 ± 626 AUs, p<0.0001, **Figure 5F-G**). We also measured contractile function in M2DCTs in each condition. We found that average baseline fractional shortening was lower in OxPhos-T3 M2DCTs (4.3 ± 1.4% vs. 5.9 ± 2.1%, 0<0.0001, **Figure 5H**). However, contraction frequency was also lower in OxPhos-T3 M2DCTs (0.31 ± 0.11 Hz vs. 0.38 ± 0.15 Hz, p=0.0001), and isoproterenol induced a much greater frequency response (2.8 ± 1.3-fold vs. 1.24 ± 0.23-fold, p<0.0001, **Figure 5I**). This greater frequency response was associated with a positive response of contractile velocity to isoproterenol in M2DCTs in OxPhos-T3 only (20 ± 0.2% increase, p<0.0001, **Figure 5J**). We also quantified differential splicing associated with maturation in *TNNT2* as an additional marker of myofilament maturation. We found that the adult *TNNT2* transcript variant accounted for 67.4 ± 0.1% of *TNNT2* transcripts in controls, 82.6 ± 1.0% of *TNNT2* transcripts in M2DCTs (p<0.0001 vs. control), and 89.9 ± 0.3% of *TNNT2* transcripts in OxPhos-T3 M2DCTs (p<0.0001 vs. M2DCTs, **Supplemental Figure 6**).

Taken together, these results indicate that the myofibrillar alignment and auxotonic contractions of M2DCTs markedly improve maturation, while exogenous metabolic and hormonal signals further drive maturation of the myofilament gene program and beta-adrenergic response.

### Controlling the Biomechanical Environment Improves Precision of Contractile Kinetic Investigation for Cardiomyopathy Disease Modeling

We next tested whether the biomechanical environment of M2DCTs would allow facile study of an hPSC-CM cardiomyopathy disease model. We compared control M2DCTs to M2DCTs generated from an isogenic line in which gene editing was performed to delete the *MYBPC3* promoter (*MYBPC3*^pr−/−^).^14^ We previously showed that this line does not express MyBP-C (encoded by *MYBPC3*), but had not quantified contractile kinetics.^14^ A complete lack of MyBP-C, as essentially occurs with homozygous truncating *MYBPC3* mutations^14^, leads to early onset, very severe hypertrophic and dilated cardiomyopathy.^29^ *MYBPC3*^pr−/−^ M2DCTs exhibited similar structural myofibrillar development as isogenic controls (**Figure 6A**). Maximal fractional shortening was reduced in *MYBPC3*^pr−/−^
 M2DCTs (4.6±1.6% vs. 5.9±2.1%, p<0.0001). Normalized contractile velocities were higher during each phase of contraction with a greater magnitude of differences in the mid and late contraction phases in *MYBPC3*^pr−/−^ M2DCTs (**Figure 6C-E,J, Supplemental Figure 7**). A more rapid contractile deceleration time was also observed in *MYBPC3*^pr−/−^ M2DCTs (**Figure 6F**). In contrast, normalized relaxation velocities were lower during each phase of relaxation in *MYBPC3*^pr−/−^ M2DCTs (**Figure 6G-I,J, Supplemental Figure 7**). These findings are consistent with the role of MyBP-C presenting a load on, and slowing myofilament velocity in the c-zone of sarcomeres^30^, such that ablation of MyBP-C results in increased contractile velocity but reduced relaxation velocity.

**Figure 6.**
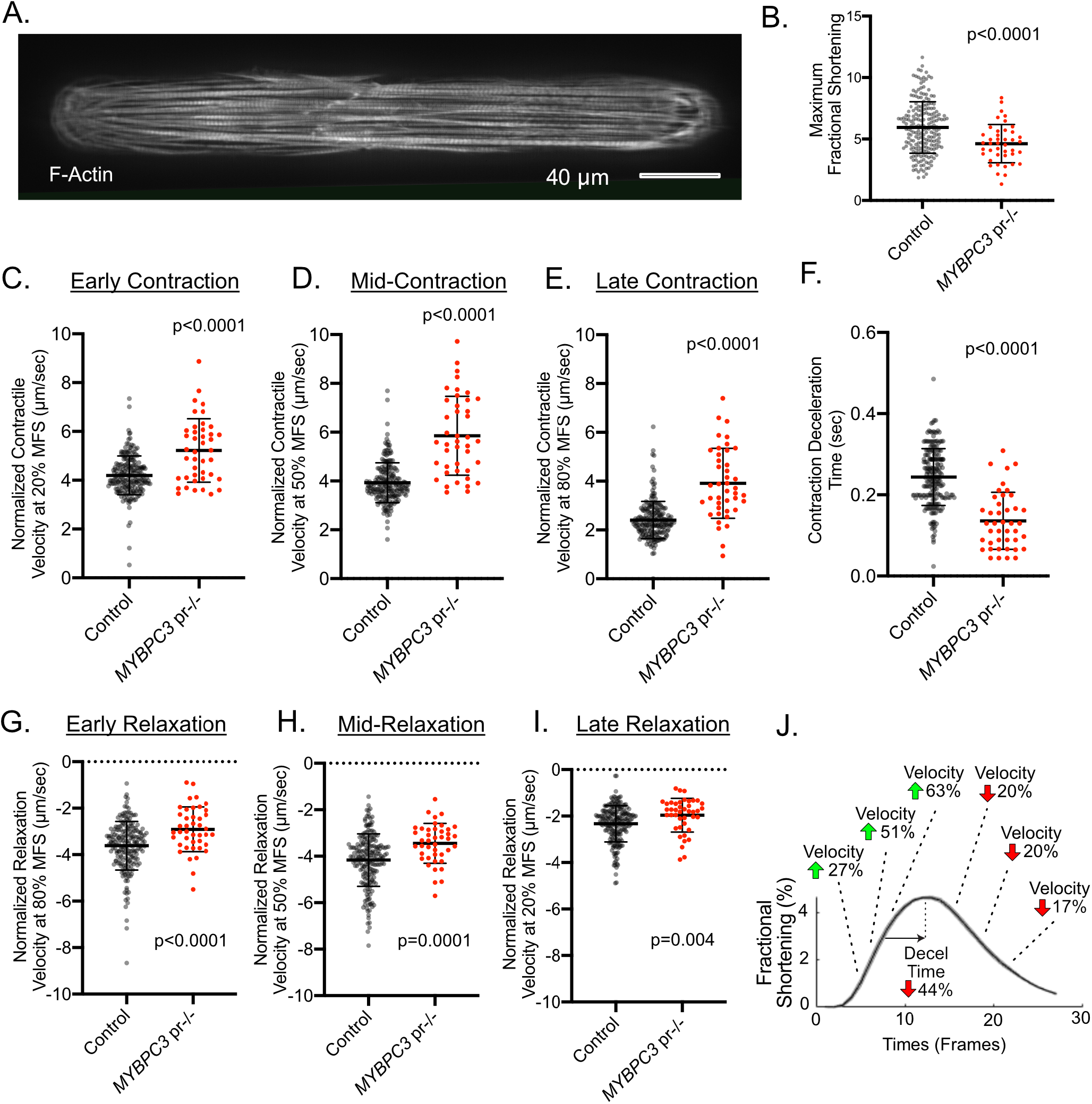
Controlling the Biomechanical Environment Improves Precision of Contractile Kinetic Investigation for Cardiomyopathy Disease Modeling. **A**. Representative example of a cardiomyopathy M2DCT (*MYBPC3*^pr−/−^) showing normally aligned myofibrils (labeled with SiR-actin). **B**. Maximal fractional shortening in control vs. *MYBPC3*^pr−/−^ M2DCTs. **C-E**. Normalized contraction velocity in control vs. *MYBPC3*^pr−/−^ M2DCTs at early (20% of max shortening), mid (50% of max shortening), and late (80% of max shortening) contraction. **F**. Contraction deceleration time in control vs. *MYBPC3*^pr−/−^ M2DCTs (defined as time from deviation from linear contractile slope to peak contraction). **G-I**. Normalized relaxation velocity in control vs. *MYBPC3*^pr−/−^ M2DCTs at early (80% of max shortening), mid (50% of max shortening), and late (20% of max shortening) relaxation. **J**. Summary of relative differences in contraction and relaxation velocities in *MYBPC3*^pr−/−^ M2DCTs shown on a representative fractional shortening curve from an *MYBPC3*^pr−/−^ M2DCT.

## Discussion

Developmental immaturity and heterogeneity, as well as technical hurdles in measuring contractile function, have been major barriers to the reproducible study of cardiac muscle derived from hPSCs in 2D.^4,5^ Although 3D constructs have been able to improve maturation, the complex 3D tissue environment, which incorporates both biomechanical cues and heterogeneous cell types, poses challenges when attempting to parse the relative importance of these distinct inputs and the technical challenges in generating state-of-the-art 3D tissues place the technique out of reach of many investigators. We created a simplified 2D tissue approach that isolates biomechanical inputs on cardiac muscle development in the absence of stromal cells. With this approach, we demonstrate that the biomechanical environment is independently a critical determinant of myofibrillar and electrophysiologic maturation in hPSC-CM tissues. Moreover, M2DCTs enable study of integrative effects from biomechanical and orthogonal biochemical signals that affect cardiac maturation. Finally, we show that rigorous control of tissue geometry and workload in M2DCTs markedly improves reproducibility of contractile quantification for pharmacologic testing and cardiomyopathy disease modeling.

Micropatterning or nanogroove techniques have been shown to generate anisotropic 2D cardiac tissues with improved myofibrillar alignment.^31,32^ However, generation of myofibrillar alignment without enabling homogeneous auxotonic contractions leads to regional tethering of 2D tissues since the entire substrate cannot be deformed commensurate with fractional shortening (one notable exception being anisotropic thin muscle films).^33^ The key insight enabling the M2DCT approach was that micropatterned cardiac tissues must be uncoupled both from each other and the rigid glass underlying the elastomer surface to allow individual cardiac tissues to experience auxotonic contractions unperturbed by neighboring cells. We designed M2DCTs to control hPSC-CM force transmission by scaling micropatterns to a size that enables 1) hPSC-CM tissue organization, 2) separation of individual tissues to avoid interactions, and 3) capability for live cell quantification. This approach generated arrays of individual tissues that rapidly align myofibrils and develop symmetric and homogeneous contractile behavior. By 10 days, these M2DCTs also exhibit further development of gap junctions and improved electrophysiology. Comparisons to standard 2D plating conditions demonstrate that myofibrillar alignment and auxotonic contractions in M2DCTs are sufficient to drive a significant level of hPSC-CM maturation even without the complex, admixed-cell environment of 3D collagen- or fibrin-based tissues.

A major motivation for this study was the variability in contractile phenotypes and extensive time required to perform contractile analyses using the traction force method in single micropatterned hPSC-CMs.^11,14^ We found that myofibrillar development was stalled in many single hPSC-CMs, with many cells unsuitable for analysis and marked variability in contractile phenotypes present both within and across differentiation batches.^11^ In contrast, multicellular connectivity and uniaxial contractions of M2DCTs were associated with robust myofibrillar development and contractile function, resulting in greater reproducibility across replicates. Throughput for contractile analysis was further increased by the ContractQuant algorithm. Whereas traction force analysis from fluorescent bead displacements requires substantial computational resources, the cross-correlation pixel tracking method in ContractQuant reduces computational time by ~10^3^-fold, enabling rapid batch processing. Moreover, brightfield imaging avoids fluorescence-induced phototoxicity as a potential experimental confounder. Using M2DCTs with ContractQuant analysis allowed precise characterization of a cardiomyopathy hPSC-CM model with contractile kinetic dysregulation due to *MYBPC3* ablation that reinforces key findings from biophysical studies.^30^ As compared to 3D microtissues or traction force microscopy of single hPSC-CMs, both the platform and analysis approaches developed here could be readily implemented in most laboratories with hPSC-CM experience.

Additionally, M2DCTs enable more facile testing of cardiac maturation strategies in a reductionist and controlled setting of purified hPSC-CMs. Our analysis showed evidence of maturation with several previously published techniques in standard 2D monolayers, but a larger gain was attained with M2DCTs, emphasizing the importance of a physiologic contractile environment for cardiomyocyte development. Superimposed on this biophysical environment, induction of oxidative phosphorylation in combination with thyroid hormone synergistically increased cTnI expression, the mature *TNNT2* transcript variant, and the beta-adrenergic response. This analysis demonstrates the utility M2DCTs to examine combinatorial influences of cardiac maturation signals on specific maturation endpoints.

In conclusion, our results show that the biomechanical environment of cardiac muscle cells derived from hPSCs is a critical determinant of their developmental maturation and contractile function. The controlled geometry and elastic workload of M2DCTs markedly improve myofibrillar, electrophysiologic, and contractile function. In addition to increasing the reproducibility of contractile assessment, M2DCTs can also serve a critical function as a testbed to study maturation in a biomechanically controlled and simplified environment deplete of stromal cells, and they enable precise contractile kinetic analyses for cardiomyopathy disease-modeling.

## Methods

### Stem Cell and Cardiomyocyte Culture

Control iPSCs were obtained from WiCell (DF19-9-11) and the Allen Institute (connexin-43-GFP, line ID = AICS-0053 cl.16). All iPSCs were cultured in StemFlex (Gibco). iPSCs were verified free of mycoplasma using MycoAlertTM (Lonza). Cardiac differentiations were performed using Wnt modulation.^1,2^ We achieved optimal differentiations using RPMI plus B27 supplement without insulin during day 0-2 (CHIR 5-6 µM during first 24 hours, LC Laboratories), then CDM3 media (without B27) with IWP4 5 µM (Stemgent) during day 3-4. Switching to CDM3 at day 3 avoids retinoic acid (included in B27), and we additionally included retinol inhibitor (BMS 453, Cayman Chemical, 1 µM) for days 3-6 to minimize atrial lineage differentiation.^35^ hPSC-CMs were maintained in CDM3 until purification by metabolic selection in CDM3-lactate using glucose-deprived, lactate-containing media on days 12-16 with one modification – we used RPMI without glutamine (Biologic Industries) to achieve a more consistent purification.^2,36^ Purified hPSC-CMs were replated as monolayers (400,000 cells/cm^2^) for 8 additional days (until day 24) on stiff PDMS membranes (Specialty Manufacturing, Inc.) coated with growth factor reduced Matrigel (Corning) in CDM3-lactate media (4 days) and then RPMI plus B27 supplement (4 days).^37^ Maintaining purified monolayers for 8 days minimized replication that occurs in earlier hPSC-CMs prior to M2DCT replating.^25^ At day 24, hPSC-CMs were dissociated by 0.25% Trypsin-EDTA (Gibco) with 5% (v/v) Liberase (Sigma-Aldrich) for 5 minutes, stopped by equal volume of 20% FBS/1 mM EDTA/PBS (maintaining low calcium at this step improves viability). The cells were triturated by gently pipetting with a p1000 pipette 3 times to obtain a near single cell suspension and then spun down at 150 cfg for 3 minutes. hPSC-CMs were resuspended in replating media (RPMI plus B27 supplement with 2% FBS) and passed through 40 um cell strainers. Live cells were counted (LUNA^™^ cell counter, typical viability ~99% with this method) and 45,000 cells were seeded per M2DCT substrate within 1.2 ml in the mini-well of devices, as described below. Two hours after cell seeding cells, the media was gently removed without drying the substrate and 4 ml of fresh replating media was added back in the dish. On the following day the media was replaced with 4 ml of 50:50 RPMI:DMEM with 1X B27 supplement. The media calcium concentration was gradually raised from 0.4 mM in RPMI to 1.1 mM by replacing half volume in the dish every 2 days with 25:75 RPMI:DMEM with 1X B27 supplement. M2DCTs were assessed after 8 additional days (i.e. day 32 post-differentiation) except where otherwise indicated. All media were equilibrated (30 minutes at 37°C and 5% CO_2_) before use.

For maturation testing, we utilized a media intended to increase oxidative phosphorylation, referred to as “OxPhos”. This media, adapted from Correia, et al., is based on glucose-free RPMI with 1X B27 supplement and galactose, lactate, glutamax, and pyruvate at final media concentrations of 4 mM, 4 mM, 2 mM, and 0.5 mM, respectively (all from Sigma).^38^ A low concentration of fatty acids is present in the B27 supplement and additional fatty acids were not added to avoid potential for oxidation and free radical formation. Triiodo-L-thyronine (T3) was used at 100 nM (Sigma), dexamethasone at 1 µM (Cayman), and torin1 at 200 nM (Cayman).

### M2DCT Substrate Fabrication

To create the M2DCT platform, micropattern geometries were first designed in AutoCAD by scaling up our prior single cell micropatterning approach.^11,14^ The design consisted of a rectangular array of 4772 rectangles with a dimension of 308 µm × 45 µm (area 13,952 µm^2^), spaced by 240 µm from each other in the rectangles’ long axis direction and by 80 µm from each other in the rectangles’ short axis direction. Standard soft lithography techniques were used to create a silicon master mold with micropattern designs from which polydimethoxysiloxane (PDMS) stamps were made. A 0.09-inch thick, 5 × 5-inch chromium photomask containing 6 replicates of each design was produced by Photo Sciences, Inc. (Torrance, CA, USA) on a soda lime glass substrate at ±1 µm feature tolerance and with the chrome side down. Microfabrication of the silicon master mold followed the standard protocol developed by Microchem, Inc. for SU-8 2005. Briefly, a 100 mm silicon wafer was first dehydrated on a hot plate at 150 °C for 5 minutes. Then a negative photoresist, SU-8 2005, was spin-coated on the silicon wafer at 500 rpm for 10 seconds and 1000 rpm for 30 seconds, which gave a thickness of ~8.6 µm. The SU-8 photoresist was then exposed to UV light at 115 mJ/cm^2^ for 9 seconds under a contact aligner (Karl Suss, MJB45) with the chromium photomask. After development of the SU-8 photoresist, the silicon wafer was hard-baked at 200 °C for 10 minutes. The thickness of the photoresist was measured using a stylus profilometer (Dektak 6M). The silicon wafer was silanized by placing in a vacuum with 2 drops of trichloro(1H,1H,2H,2H-perfluorooctyl)silane placed on aluminum foil. To prepare the PDMS stamps, Sylgard-184 elastomer and curing agents (Dow Corning, Midland, MI) were mixed at a ratio of 10:1 (w/w), degassed, then cast over the silicon mold and cured at 50 °C overnight. Individual stamps were cut from the cured PDMS. Individual stamps can be re-used innumerable times as long as the surface is not inadvertently scratched.

Initial micropatterning testing was performed on polyacrylamide hydrogels, fabricated onto glass coverslips and microprinted with fibronectin (Sigma) as previously described.^14,16^ Since this method failed to maintain adherence of M2DCTs, final optimized M2DCT substrates consisted of micropatterned 8 kPa PDMS. Soft PDMS at specified stiffness was formulated by mixing Sylgard® 527 and Sylgard® 184 as described.^17^ Specifically, each component was first mixed with its own curing agent (i.e. 50:50 for Sylgard® 527 and 10:1 for Sylgard® 184). To obtain 8 kPa PDMS, a 1:97 ratio (mass:mass) of Sylgard 184 : Sylgard 527 was mixed thoroughly, then degassed.

Glass bottom dishes were fabricated by first gluing a 40 mm diameter glass coverslip underneath a 6 cm culture dish with a 25 mm hole drilled through the bottom using silicon all-purpose adhesive sealant (DAP), creating a 25 mm mini-well in the center of the 6-cm dish. 135 µl of the PDMS mixture above was pipetted onto the center of each mini-well with a P1000 pipette and low-viscosity tip to create a ~70 µm thick layer of PDMS. Since 8 kPa PDMS is too tacky for direct microcontact printing, a 2-step transfer process with a fibronectin micropatterned polyvinyl alcohol (PVA) film intermediary was used.^18,19^ To create the PVA film, 65 ml of 5% (w/v), 70 µm strainer filtered, PVA solution was poured on to a 20.5 cm × 25.5 cm leveled flat glass panel and left to air dry (2-3 days). Completely dried PVA films were peeled and trimmed. To print micropatterns, PDMS micropatterning stamps were first cleaned in 50% ethanol sonication bath. 350 μl of 1:25 diluted 0.1% human serum derived fibronectin (Sigma-Aldrich) was added per stamp and incubated at room temperature for 1 hour, then aspirated. Following air drying, stamps were pressed on pre-cut, individual stamp-size PVA films and incubated for 30 minutes. The PVA films with fibronectin micropatterns were then peeled and inverted onto cured 8 kPa PDMS. Microprinted dishes were stored up to 2 weeks at 4°C prior to use with the PVA films remaining adherent. Immediately prior to use, PVA films were dissolved in sterile water at room temperature on an orbital shaker (70 RPM) for 15 minutes to expose the micropattern, and surfaces were sterilized by UV in the culture hood for 10 min.

### Microscopy

Microscopy was performed at 37°C and 5% CO_2_ for live cell experiments. DIC microscopy was performed using a Nikon Eclipse Ti-E inverted microscope with motorized stage and an Andor Zyla camera. M2DCTs filling micropatterns with approximately 6-12 cells per pattern were first identified at 10X and locations saved to increase efficiency of imaging. A 40x air gap objective was then positioned for contractile imaging (avoiding heat sink from oil immersion) and time-series images were obtained for each M2DCT with field of view 330 × 100 µm at 50 frames per second. All hPSC-CMs were labeled for F-actin with the cell permeant dye, SiR-Actin (Cytoskeleton, Inc.) prior to imaging (0.5 µM, 1 hour) to enable visualization of myofibrils following contractile imaging, and this protocol enabled confirmation that all selected M2DCTs were 100% pure cardiomyocytes in every experiment. Using the tagged locations of M2DCTs previously imaged for contractions, F-actin images were obtained as a z-stack in fixed (4% paraformaldehyde) M2DCTs with 1 µm slice thickness, and deconvolutions with maximum intensity projections were performed in ImageJ.

Time lapse imaging of the connexin-43-GFP reporter line was performed with confocal microscopy at 40X. For the connexin-43 imaging only, myk-461 was first administered at 500 nM prior to imaging to temporarily reduce fractional shortening to minimize blurring during z-stack acquisitions with 0.5 µm slice thickness. Image locations were saved during imaging at day 3, and the same tissues were imaged again at day 10. As above, SiR-Actin staining was performed prior to both day 3 and day 10 for concurrent imaging of myofibrils. Maximum intensity projection images were obtained from each z-stack. Myofibrillar orientation images were obtained by labeling fixed (4% paraformaldehyde) M2DCTs with an MyBP-C antibody (N-terminal rabbit polyclonal, 1:1000, kind gift from Samantha Harris) and anti-rabbit AlexaFluor secondary (ThermoFisher). Intercalated disk images were obtained by labeling fixed M2DCTs with an N-cadherin antibody (1:200, BD Biosciences) and anti-mouse AlexaFluor secondary (ThermoFisher).

### Image Processing

Contractile function was quantified from DIC images using a pixel cross correlation method that we implemented in MATLAB, referred to as ContractQuant. ContractQuant tracks displacement of M2DCTs by solving the best match for regions of interest (ROIs) placed at the 50% inner length of each M2DCT. ROIs can be modified or manually selected for different imaging platforms. ContractQuant exports the following measurements: 1) contractile velocities at 20%, 50%, and 80% of peak contraction; 2) relaxation velocities at 20%, 50%, and 80% of peak contraction; 3) max fractional shortening; 4) contraction acceleration; 5) contraction deceleration time; 6) relaxation time. Standard deviations for each parameter and graphs of displacement, velocity, acceleration, and a symmetrical motion index are also exported, enabling rapid confirmation of accurate tissue tracking. Rare cases with inconsistent tissue tracking were excluded from analysis.

Individual sarcomere unit orientations were measured across images using a separate automated imaging method in MATLAB that creates a binary mask of segmented sarcomere units. These orientations were fit to a Gaussian distribution for each image to calculate the standard deviation of sarcomere angles per image. Myofibrillar abundance in M2DCTs was measured using a previously described MATLAB script that detects myofibrils traversing the long-axis of tissues based on F-actin signal and then quantifies density and heterogeneity of these structures.^11^

### Intracellular Calcium Quantification

Cardiomyocytes were loaded with 1 µM fura-2 AM (Thermo Fisher) for 10 min, rinsed with 250 µM probenecid in HBSS for 5 min, changed to pre-equilibrated media, and then incubated for 30 min prior to imaging through a Nikon Super Fluor 40x objective. The background-subtracted emitted light intensity following alternating excitation at 340 nm and 380 nm was reported as fura-2 ratios.

### Patch Clamping

Whole-cell patch clamp was performed in current-clamp mode using an Axopatch 700B amplifier and Digidata 1440A (both from Axon Instruments) at room temperature. M2DCTs were bathed in a solution containing (in mM): 135 NaCl, 4 KCl, 1.8 CaCl_2_, 1 MgCl_2_, 10 Hepes, 1.2 NaH_2_PO_4_, 10 glucose, pH 7.35 with NaOH. Patch pipettes (2-3 MΩ) were filled with the internal solution containing (in mM): 130 K-aspartate, 10 KCl, 9 NaCl, 0.33 MgCl_2_, 5 Mg-ATP, 0.1 GTP, 10 Hepes, 10 glucose, pH 7.2 with KOH. The resting membrane potential (RMP) was determined under current clamp at zero current. Action potentials (APs) were elicited by 20 pulses of 0.4 nA each at 0.5 Hz. The liquid junction potential between bath solution and pipette solution was calculated with pClamp and corrected after experiments.

### RNA-Seq

RNA-Seq was performed on 5 pooled batches of hPSC-CMs with verified >95% cardiomyocyte purity. These lactate-purified batches had previously been cryopreserved at day 16 to enable pooling of batches together in a single thaw to enable comparison of experimental conditions independent from potential batch effect. Each condition was performed in triplicate and cultured for 8 days prior to collection. RNA samples were processed in a single session using the RNAeasy kit (Qiagen). All samples used for RNA library preparation had an RNA abundance >25 ng/µl, 260/280 ratio >1.9 (by NanoDrop^™^), and RNA integrity number >8.0. A stranded and barcoded RNA library was prepared and sequenced using a multiplexed library on an Illumina NovaSeq by the University of Michigan DNA Sequencing Core. Reads were aligned to the genome using STAR.^39^ Lowly expressed genes were filtered, leaving a dataset of 20,220 genes. Reads per gene were normalized to the library size for each sample using DESeq2.^40^ Relative abundance for each gene was determined using dispersion estimates for each gene fit to a binomial generalized linear model with DE-Seq2.^40^ The Wald test with multiple testing correction for 20,220 genes tested was performed to determine statistical significance (adjusted p-value <0.001).^40^ GOseq was used to test for enrichment of biologic pathways among differentially regulated genes with multiple testing correction for all pathways tested.^41^ These data were deposited in NCBI’s Gene Expression Omnibus (GEO accession pending).

### Mass Spectroscopy

The effect of OxPhos media on relative cTni and ssTnI abundance was quantified using mass spectroscopy to quantify unique peptides in both cTnI and ssTnI. This data was analyzed from a previously described experiment, in which pooled hPSC-CMs from 3 separate lines (DF19-9-11 and 2 *MYBPC3* patient lines, kind gift from Christine Mummery) were switched at time 0 to OxPhos media.^14^ Abundance of cTnI and ssTnI were calculated at each time point using the 3 most abundant unique peptides from each protein. The relative proportion at each time point was calculated as cTni/(cTnI+ssTnI) and ssTnI/(cTnI+ssTnI) respectively.

### Statistical Analysis

Continuous variables were compared by 2-sample t-tests or ANOVA with multiple testing correction for normally distributed outcomes. For non-normally distributed outcomes, the Mann-Whitney test was used. Serial measurements in the same tissues were compared with the paired samples t-test. Statistical tests and graphs were generated using GraphPad Prism. RNA-seq statistical analysis is described above.

## Acknowledgements

This work was funded by the NIH (K08HL130455 to ASH, R21GM134167 to APL, R01HL149363 to LLI), NSF (CMMI-1561794 to APL), Michigan Institute for Clinical and Health Research Postdoctoral Translational Science Program fellowship from UL1TR002240 to Y-TZ, and the University of Michigan Research Stimulus Award (ASH and APL).

## Author Contributions

Conceptualization - ASH, APL; Methodology – Y-CT, Y-TZ, AC, SJD, YWW, MJP, BMB, DN, LLI, APL, ASH; Investigation – Y-CT, Y-TZ, AC, SJD, BE, IP, SF, TSO, NW, KH; Writing – Y-CT, AC, ASH; Writing, Review/Editing – Y-CT, SJD, MJP, BMB, DN, LLI, APL, ASH; Funding – Y-TZ, LLI, APL, ASH; Supervision – MJP, BMB, DN, LLI, APL, ASH.

