## Supplemental Data for "Biomechanics and Myofibrillar Alignment Enhance Contractile Development and Reproducibility in Stem Cell Derived Cardiac Muscle"

**Supplemental Material**

### Supplemental Notes

<u>Modeling Analysis of M2DCT Biomechanics on Elastic PDMS Substrates</u>

A mechanics problem was set up to investigate the relationship between fractional shortening and force exerted by M2DCTs on elastomeric substrates composed of PDMS. In particular, we aimed to assess whether uncertainty in the mechanical characteristics of the M2DCT or variation in the dimensions of the PDMS influenced the relationship between fractional shortening and exerted force. The simulation was based on a single M2DCT adhered onto a PDMS substrate. To replicate the conditions of the experiments as closely as possible, the M2DCT was defined as a thin cuboid of 7:1 length to width aspect ratio - 308 µm length, 44 µm width, and 10 µm thickness. The buffering space between micropatterns was 120 µm in the long-axis direction and 80 µm in the short axis direction – corresponding to a PDMS:M2DCT ratio of approximately 1.4:1 length ratio and 2.8:1 width ratio, leading to 431.2 µm length and 123.2 µm width for the PDMS. The PDMS thickness was set to 70 µm to match manual measurements of PDMS from fabricated devices.

The material properties of the PDMS were defined using a neo-Hookean constitutive model with stiffness 8 kPa.[1] Cardiomyocyte's stiffness can vary between 2 and 12 kPa, so here the M2DCT was modelled based on an underlying neo-Hookean model with stiffness 8 kPa. Fibers were prescribed along the length of the M2DCT, which would lead to a contractility of up to 300 nN/µm$^2$ (mN/mm$^2$). This covers the range of typical myocardium contractility (~50 mN/mm$^2$). More complex passive cardiac mechanics models were feasible, but not considered here due to a lack of passive mechanical data and the amplification of the passive parameter space.

For modelling purposes, structural symmetry was assumed. Thus, only ¼ of the problem was simulated (½ of the length and ½ of the width), and the results were reflected in order to visualize the entire domain (Figure 2A). For the PDMS, a hexahedral mesh with 500 elements and 4851 quadrilateral nodes was created. For the M2DCT, a hexahedral mesh with 50 elements and 651 quadrilateral nodes was used. For the latter, some of the elements were modified into collapsed elements, as the short side of the M2DCT was modelled as tapering into the PDMS substrate, as can be seen in Figure 2A. On the bottom and sides of the PDMS, 0 displacement boundary conditions were imposed, whereas at the reflection planes, in both the PDMS and the M2DCT, no penetration was allowed. The displacements at the contact interface between the M2DCT and PDMS were restricted to be the same, i.e. the M2DCT could not slide on top of the PDMS. The force exerted by the M2DCT on the PDMS substrate was computed over the interface surface. All simulations were run in CHeart.

Simulations were run to examine how the observed fractional shortening correlates with force, as well as to understand how varying the testing conditions might affect the observations. The effect of the PDMS buffering space on the fractional shortening was investigated by varying the PDMS:M2DCT length ratio between 1.2:1 to 2:1 and the width ratio was varied between 2:1 to 3:1, both with an increment of 0.2:1. Subsequently, the effect of PDMS thickness (varying between 20 and 100 µm) on fractional shortening was examined.

Finally, the stiffness of the M2DCT was altered between 2 to 12 kPa, respectively, to study how the force vs fractional shortening behavior changes.

Although the mean fractional shortening observed in experiments was approximately 5%, the range extended to ~11%. Therefore, our simulations were performed up to a fractional shortening of ~11% to verify fidelity of the system across the entire range of experimental measurements. In Supplemental Figure 2, it can be seen that prescribing the PDMS buffer used in the experiments (1.4:1 and 2.8:1 PDMS:M2DCT length and width ratios, respectively) leads to 11.2% fractional shortening (~17.2 µm total displacement, i.e. ~8.6 µm at each end) and 11.4 µN force exerted by the M2DCT. Increasing the buffer in either length or width would lead to a similar fractional shortening reading (11.4%, an increase of 0.2%), while decreasing the buffer space would also lead to similar values (e.g. 10.7%, a decrease of 0.5%). The force varies by less than 10% between the tightest and largest buffering spaces (12.1 to 11.1 µN). When the buffer space is small, the M2DCT contracts less and exerts more force on the PDMS, compared to a larger buffer space. However, the absolute reading differences would be even less significant when the typical maximum fractional shortenings observed are ~5%. Thus, it can be inferred that the buffer space used in the experiments does not alter the fractional shortening and forces, and adjacent M2DCTs are not likely to significantly influence each other.

The next test investigated the influence of PDMS thickness on relationship between force and fractional shortening. We calculated this relationship for 20, 30, 40, 70 and 100 µm PDMS thickness (Supplemental Figure 2). A thin PDMS layer, of only 20 or 30 µm, would lead to a biased relationship, whereas it can be seen that for PDMS thicknesses larger than 40 µm the behavior does not change significantly. Thus, the experimental substrate thickness (~70 µm) does not induce a bias on the behavior of force with fractional shortening, even if a fabrication error resulted in up to a 30 µM difference.

We selected 8 kPa as the elastic modulus of M2DCTs to model the above conditions, based on prior measurements in cardiomyocytes. As previously discussed, the fractional shortening of M2DCTs was computed based on displacement readings on top of each M2DCT. Since the actual elastic modulus may vary across individual M2DCTs, we tested how varying the elastic modulus might introduce error into calculations of force. As shown in Supplemental Figure 2, a lower M2DCT elastic modulus would result in a higher measured fractional shortening due to greater deformation of the tissue relative to the stiffer 8 kPa PDMS. In contrast, the behavior of M2DCTs with elastic moduli in the range of 8-12 kPa tends towards an asymptotic behavior. This analysis shows that variability in elastic moduli across individual M2DCTs in the range of 8-12 kPa will have a negligible effect on fractional shortening and force generation on 8 kPa PDMS, while tissue elastic moduli <6 kPa will result in systematic overestimation of traction forces.

Additionally, we analyzed how ROI placement for fractional shortening measurements may affect reliability of measurements. Because of edge tapering, the relative fractional shortening does not increase linearly near the M2DCT long-axis ends (Supplemental Figure 3D). Based on analysis of fractional shortening and force relationships, we selected placement of ROIs at the inner 50% of M2DCT tissue length to minimize effects of ROI placement on measurements of contractility, since fractional shortening measurements are relatively constant in these regions (Figure 2A, Supplemental Figure 3E-F).

Finally, we generated a MATLAB function to correlate fractional shortening and traction force based on the above results with inputs of PDMS stiffness and ROI location. Example correlations are shown in Table 1 summarizes the correspondence between fractional shortening and force for the case of M2DCTs on 70 um thick, 8 kPa PDMS.

**Supplemental Table 1. Gene Ontology – Biologic Process All Results**

**Supplemental Table 2. Gene Ontology – Biologic Process Summary Results**

**Supplemental Video 1.** Representative video of an M2DCT from brightfield imaging at 40X. M2DCTs typically exhibit spontaneous contractions with symmetric and homogeneous contractions in the uniaxial direction of the M2DCT long axis.

**Supplemental Video 3.** Live imaging of myofibrils from standard (nonpatterned) iPSC-CMs. Myofibrils were marked for F-actin (SiR-actin) for live-cell imaging. Regional heterogeneity in magnitude of contractions causes some regions to experience myofibrillar stretch due electromechanical coupling to adjacent, stronger-contracting cells.

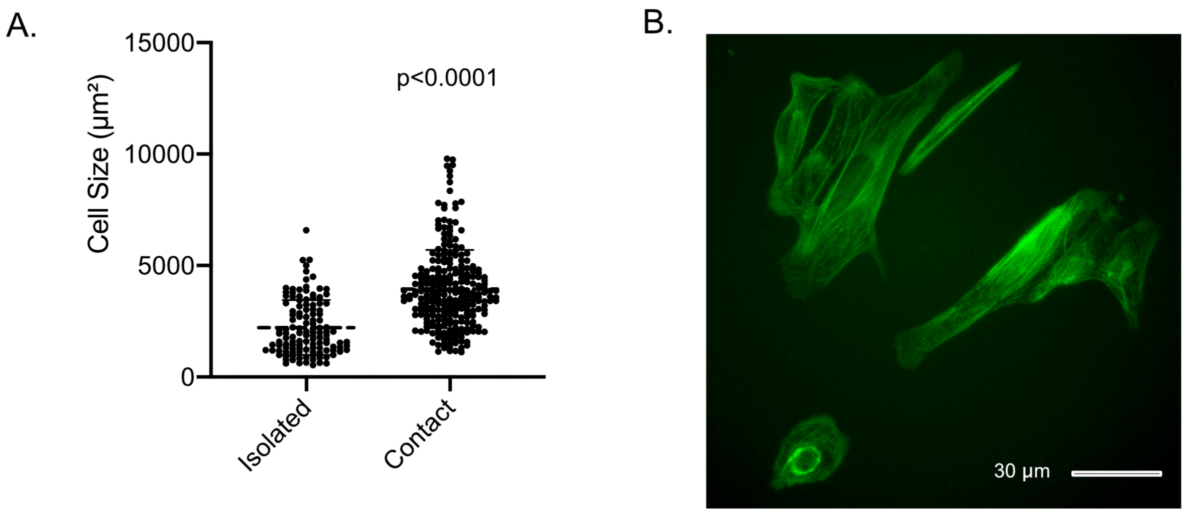


**Supplemental Figure 1. Cell contact drives iPSC-CM growth. A.** Cell area of iPSC-CMs was measured from phalloidin-stained images of iPSC-CMs plated at intermediate density to control for effects of batch or media conditions. Cell area was analyzed to compare sizes of isolated single cells versus cells in contact with at least one other cell. **B.** Representative image showing cell sizes in isolated iPSC-CMs compared to those in direct contact with other iPSC-CMs.

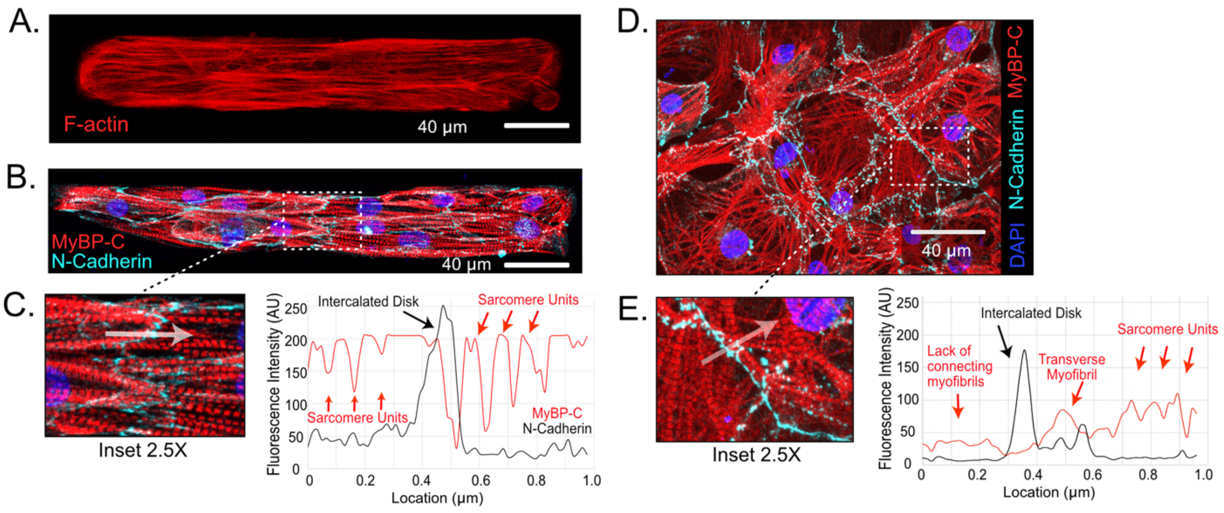


**Supplemental Figure 2. Myofibrillar alignment rapidly develops in M2DCTs and is associated with organization of intercellular junctions. A.** Representative example of M2DCT showing myofibrillar alignment by 3 days following dissociation and re-plating (stained by SirActin to label F-actin). **B.** Representative M2DCT showing myofibrillar organization at cell junctions by the intercalated disk protein, N-cadherin. **C.** Inset (2.5X) and intensity profile (measured across opaque arrow) shows that myofibrillar bundles are aligned at cell junction connection points with continuation of sarcomeric periodicity across the border junction. **D.** Representative standard iPSC-CMs, showing myofibrillar disorganization within and between cells that prevents consistent myofibrillar alignment across cell junctions. **E.** Inset (2.5X) and intensity profile (measured across opaque arrow) shows that myofibrillar bundles are not consistently aligned at cell junction connection points in standard iPSC-CMs. In the region measured with the intensity profile, no aligned myofibrils are present left of the junction. Right of the junction, a myofibrillar bundle runs parallel to the junction, which is transverse to other myofibrillar bundles connecting to the junction (the perpendicular myofibrils exhibit sarcomeric periodicity along the profile).

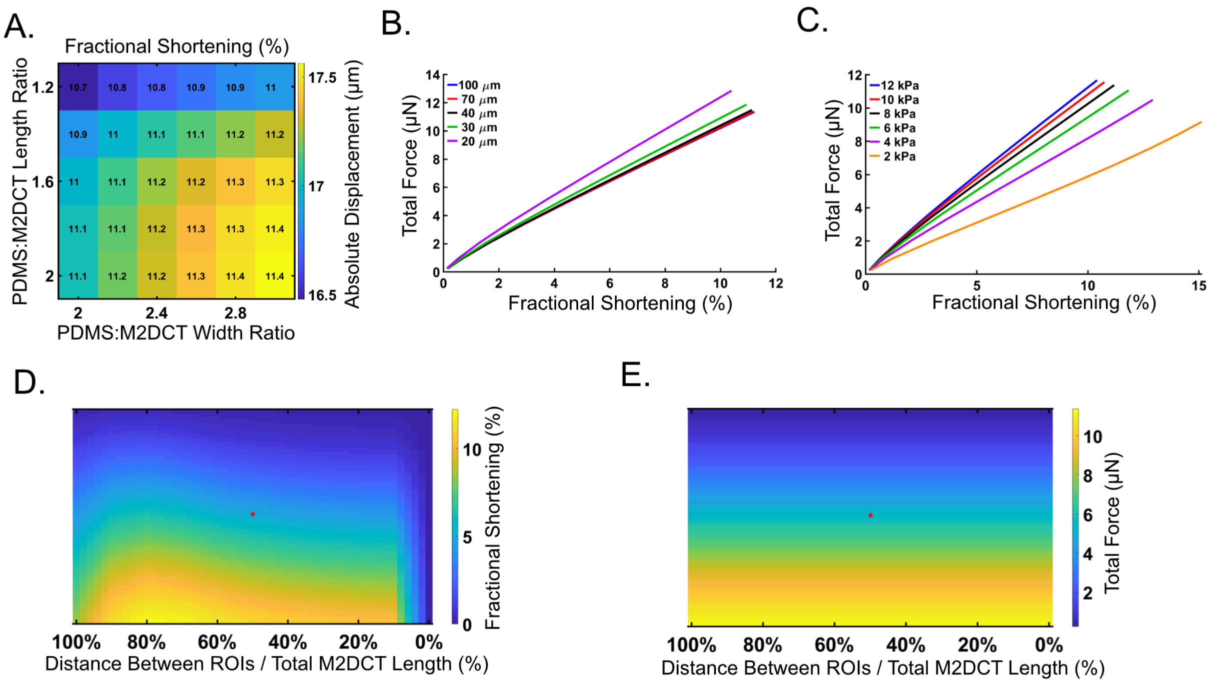


**Supplemental Figure 3. Modeling shows minimal boundary effect and minimal impact of substrate thickness on M2DCT force development while enabling conversion of ROI displacements to whole tissue force. A.** The M2DCT substrate design includes a buffering region of PDMS surrounding each tissue with a ratio of 1.4 of PDMS:M2DCT along the tissue length and 2.8 along the tissue width. Modeling the deformation of 8 kPa PDMS by an M2DCT with 11% fractional shortening (>95$^{th}$ percentile of observed fractional shortening) demonstrates that further increasing the buffering region resulted in only minor changes in the relationship between force and tissue displacement. **B.** Variation in PDMS thickness in the range of 40-100 µm demonstrates negligible effects on the fractional shortening and force relationship, while 30 µm thickness and lower requires greater force to achieve a given fractional shortening due to tethering from the underlying glass. **C.** The effect of tissue elastic modulus on underlying PDMS deformation and force generation at given fractional shortening magnitudes was assessed by modeling each using a neo-Hookean model. In the range of cardiac tissue elastic moduli (8-12 kPa), the influence of variability in M2DCTs' elastic moduli was minor on 8 kPa PDMS and relationships were approximately linear. **D-E.** The effect of region of interest placement for tracking fractional shortening was evaluated by modeling the dispersion of regional displacements within an M2DCT to attain a given whole tissue force assuming homogeneous distribution of contracting myofibrils within the tissue and tapering of tissue edges at the lengthwise ends. D and E show corresponding fractional shortening (D) and whole tissue force (E) heat maps with local correlations as a function of distance between the measured ROIs on the x-axis. Near the lengthwise tissue edges (100%) and near the tissue center (0%), small differences in ROI placement exert a large influence on the correlation between fractional shortening and whole tissue force. We selected ROI placement to capture displacements of the inner 50% of M2DCTs since this location is robust to minor errors in ROI location (matching red asterisk in the center of each heat map corresponds to inner 50% ROI measurements of an M2DCT contracting with 5% fractional shortening to generate 5.48 µN of total force. The relationships shown in D-E are incorporated into a MATLAB script with a tabulated look-up table that calculates whole tissue force as a function of PDMS stiffness, fractional shortening, and ROI location.

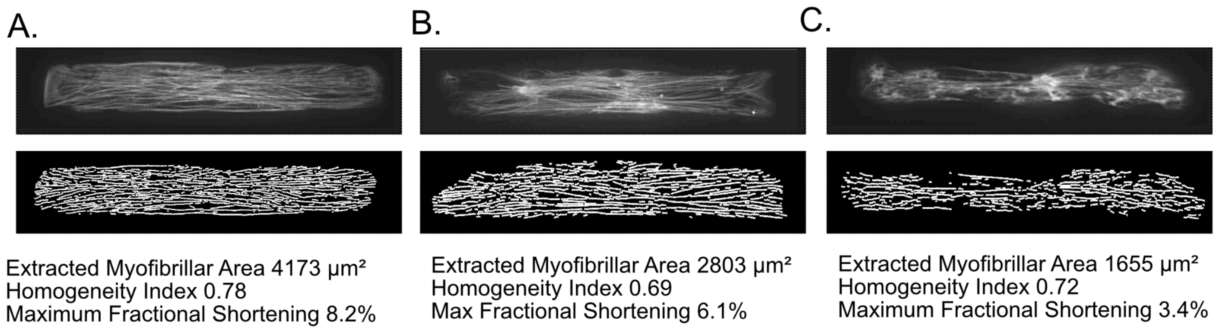


**Supplemental Figure 4. Myofibrillar abundance correlates with differences in contractile function. A-C.** Representative examples of M2DCTs with varying myofibrillar structural development. Myofibrils were imaged by labeling F-actin (SiR-actin) immediately following contractile imaging and then obtaining z-stacks of images. Automated analysis of deconvoluted images was performed in MATLAB to extract signal peaks from myofibrils, allowing calculation of total myofibrillar bundle area and an index of homogeneity of myofibrils within the M2DCTs. Greater myofibrillar area (as in A) generally correlated with larger magnitudes of fractional shortening. Poorly formed myofibrils (as in C) were present in a low proportion (<5%) of imaged M2DCTs, and these M2DCTs were excluded from analysis to avoid bias due to batch variability.

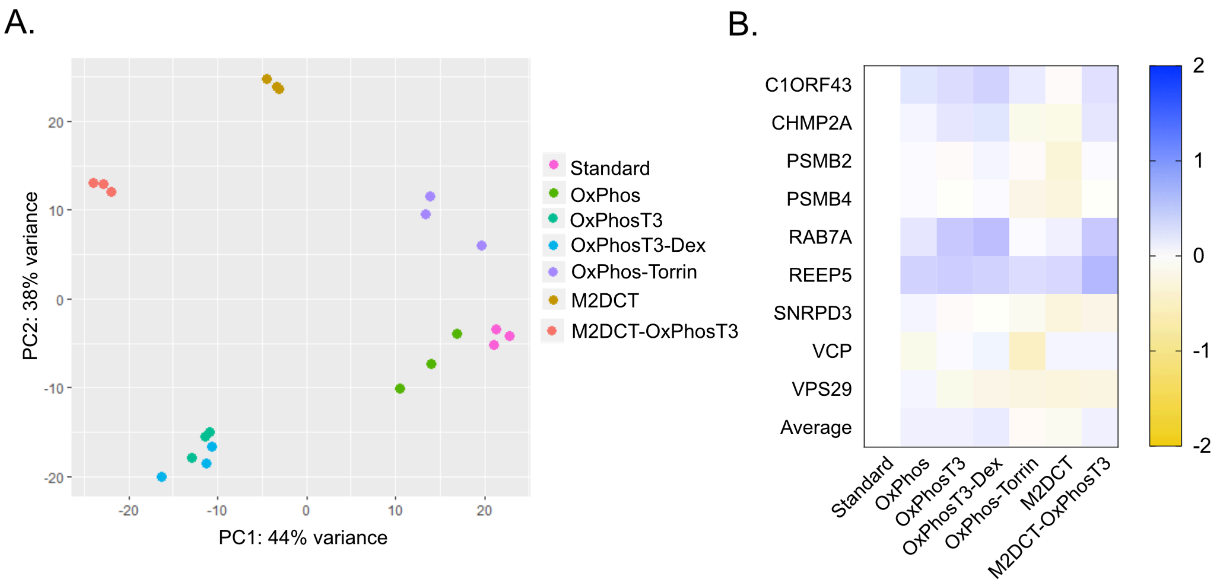


**Supplemental Figure 5. Principal component analysis reveals clustering between samples in different maturation conditions while housekeeping gene expression was similar. A.** Principal component analysis of normalized RNA-seq reads across different maturation conditions shows independent clustering of biologic replicates (N=3) for each condition except for similar clustering among OxPhosT3 and OxPhosT3-Dex groups. **B.** Relative expression (log-2 fold chance) of a panel of housekeeping genes across maturation conditions shows similar expression profiles indicating robust normalization of RNA-seq reads by DE-Seq2.

A.

B.

C.

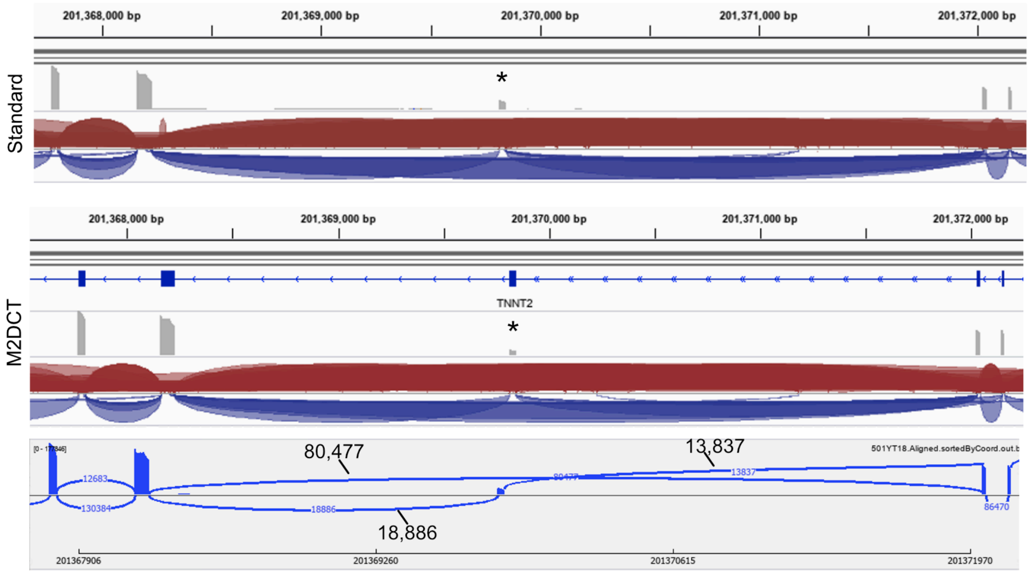


**Supplemental Figure 6. M2DCTs exhibit a larger proportion of splice exclusion of exon 5 of TNNT2. A.** Read depth of RNA-seq reads for *TNNT2* surrounding exon 5 is shown for standard iPSC-CMs for a representative sample, shown from the IGV genome browser in the top track. In standard iPSC-CMs, 33% of TNNT2 transcripts contained exon 5 (PSI 32.6 ± 0.1%) indicating that 67% of transcripts are the adult variant (exon 5 marked by *). The lower track graphically depicts spice junction reads that include or exclude exon 5. **B.** In M2DCTs, greater exclusion of exon 5 was present (PSI 17.4 ± 1.0%, p<0.0001) indicating that 83% of transcripts are the adult variant. **C.** A Sashimi plot shows numbers of reads spanning splice junctions that either include or exclude exon 5 from a representative M2DCT.

A. B.

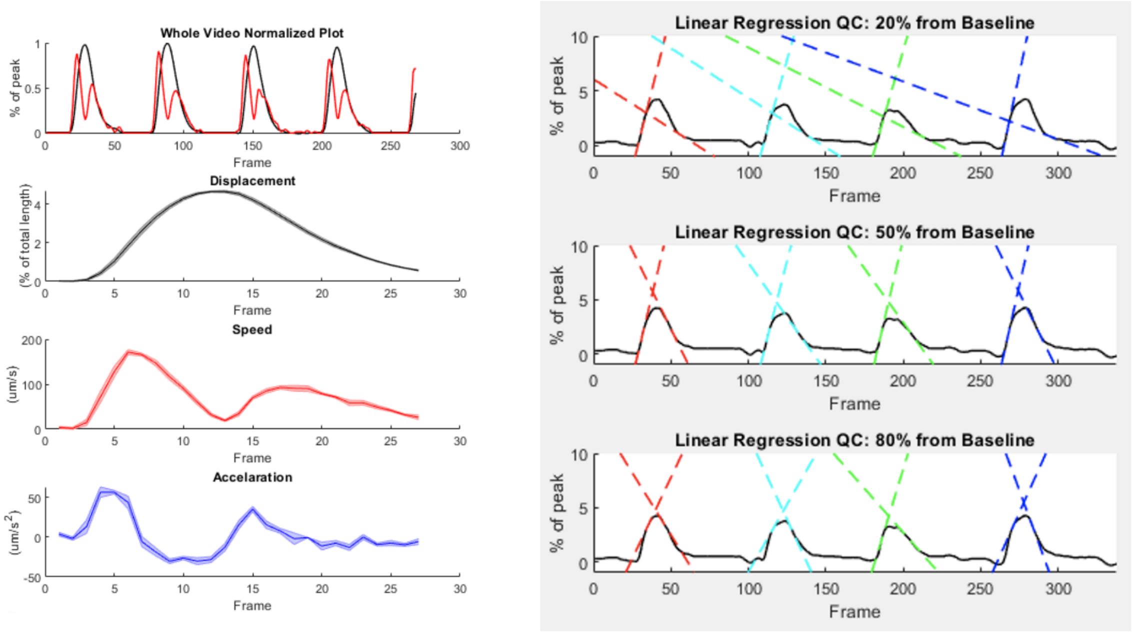


**Supplemental Figure 7. ContractQuant enables kinetic analysis of *MYBPC3*$^{pr-/-}$ M2DCTs. A.** Example output from ContractQuant from an *MYBPC3*$^{pr-/-}$ M2DCT showing normalized displacement (black) and velocity (red, top row), fractional shortening (second row), velocity (third row), and acceleration (bottom row). The median value for each parameter is extracted for subsequent analyses. The merged contraction graphs ($2^{nd}$-$4^{th}$ rows) generated by ContractQuant enable rapid verification of consistent tissue tracking and symmetric tissue contraction. **B.** Example output from ContractQuant showing analysis of contraction and relaxation velocities from an *MYBPC3*$^{pr-/-}$ M2DCT at 20% (top), 50% (middle), and 80% (bottom) of peak contraction. The median value is extracted for subsequent analyses. The figures generated by ContractQuant enable rapid assessment of accurate automated identification of early, mid, and late time points for velocity calculations for each tissue.
